# Dietary salt promotes cognitive impairment through tau phosphorylation

**DOI:** 10.1101/470666

**Authors:** Giuseppe Faraco, Karin Hochrainer, Steven G. Segarra, Samantha Schaeffer, Monica M. Santisteban, Ajay Menon, Hong Jiang, David M. Holtzman, Josef Anrather, Costantino Iadecola

## Abstract

Dietary habits and vascular risk factors promote both Alzheimer’s disease and cognitive impairment caused by vascular factors^1-3^. Furthermore, accumulation of hyperphosphorylated tau, a microtubule associated protein and a hallmark of Alzheimer’s pathology, is also linked to vascular cognitive impairment^4-7^. In mice, a salt-rich diet leads to cognitive dysfunction associated with a nitric oxide deficit in cerebral endothelial cells and cerebral hypoperfusion^8^. Here we report that dietary salt induces tau hyperphosphorylation followed by cognitive dysfunction, effects prevented by restoring endothelial nitric oxide production. The nitric oxide deficiency reduces neuronal calpain nitrosylation resulting in enzyme activation, which, in turn, leads to tau phosphorylation by activating cyclin dependent kinase-5. Salt-induced cognitive impairment is not observed in tau-null mice or in mice treated with anti-tau antibodies, despite persistent cerebral hypoperfusion and neurovascular dysfunction. These findings unveil a causal link between dietary salt, endothelial dysfunction and tau pathology, independent of hemodynamic insufficiency. Avoiding excessive salt intake and maintaining vascular health may help stave off vascular and neurodegenerative pathologies underlying late-life dementia.

---

Vascular risk factors including excessive salt consumption have long been associated with cerebrovascular dysfunction and cognitive impairment^1-3^. A diet rich in salt promotes stroke and dementia independently of hypertension^9-12^ and has been linked to the cerebral small vessel disease underlying vascular cognitive impairment^13,14^, a condition associated with endothelial dysfunction and reduced cerebral blood (CBF)^15^.

Accumulation of the microtubule associated protein tau is a pathological hallmark of Alzheimer’s disease^16^. Excessive tau phosphorylation promotes the formation of insoluble tau species, thought to mediate neuronal dysfunction and cognitive impairment^17,18^. Tau hyperphosphorylation and aggregation have increasingly been detected also in vascular brain pathologies both in humans and animal models^4-6^, and have been linked to cognitive dysfunction in patients with small vessel disease^7^.

In mice, a high salt diet (HSD) induces cognitive dysfunction by targeting the cerebral microvasculature through a gut-initiated adaptive immune response mediated by Th17 lymphocytes^8^. The resulting increase in circulating IL17 leads to inhibition of endothelial nitric oxide (NO) synthase (eNOS) and reduced NO production in cerebral microvessels, which, in turn, impairs the endothelial regulation of microvascular flow and lowers cerebral blood flow (CBF) by ≈25%^8^. Remarkably, the increases in CBF evoked by neural activity and blood-brain permeability are not altered^8^. However, it remains unclear how hypoperfusion, in HSD as in other vascular risk factors, leads to impaired cognition. The prevailing view is that reduced CBF compromises the delivery of oxygen and glucose to energy-demanding brain regions involved in cognitive function^15,19^. But the relatively-small reductions in CBF associated with HSD^8^ and vascular cognitive impairment^20^ do not reach the threshold needed to induce sustained cognitive dysfunction (≥50% CBF reduction)^21,22^. Thus, vascular factors beyond cerebral perfusion could also be involved.

To address this question, we investigated whether tau contributes to the cognitive impairment induced by HSD and, if so, whether the effect depends on the associated cerebral hypoperfusion. First, we established if HSD induces tau phosphorylation. Male C56Bl/6 mice were placed on a normal diet (ND) or HSD (4 or 8% NaCl), corresponding to a 8-16 fold increase over the salt content in the regular mouse chow and approaching the highest levels of human salt consumption^23^. Phosphorylation of selected tau epitopes linked to tau aggregation and neuronal dysfunction^17^ were assessed 4, 8, 12, and 24 weeks later by Western blotting. HSD (8%) increased p-tau (AT8, RZ3 epitopes) in neocortex and hippocampus without upregulation of total tau (Tau 46) (Fig. 1A). In the hippocampus, an increase in PHF13 was also observed (Extended Data Fig. 1A). HSD did not increase tau acetylation, a post translational modification implicated in the harmful neuronal effects of tau^24^ (Extended data Fig. 1A). AT8 and RZ3 were also increased in neocortex of female mice fed a HSD (Extended data Fig. 1B). AT8 and MC1 immunoreactivity was detected in neuronal cell bodies of the pyriform cortex and other cortical regions, but neurofibrillary tangles were not observed (Fig. 1B-C; Extended Data Fig. 1C-D). As anticipated, p-tau (AT8) was abolished by incubation of the sample with lambda protein phosphatase (Extended Data Fig. 1E). Increased AT8 was also observed in neocortex with a 4% HSD (Extended Data Fig. 1F), indicating that also lower amounts of dietary salt are sufficient to induce tau phosphorylation.

**Figure 1.**
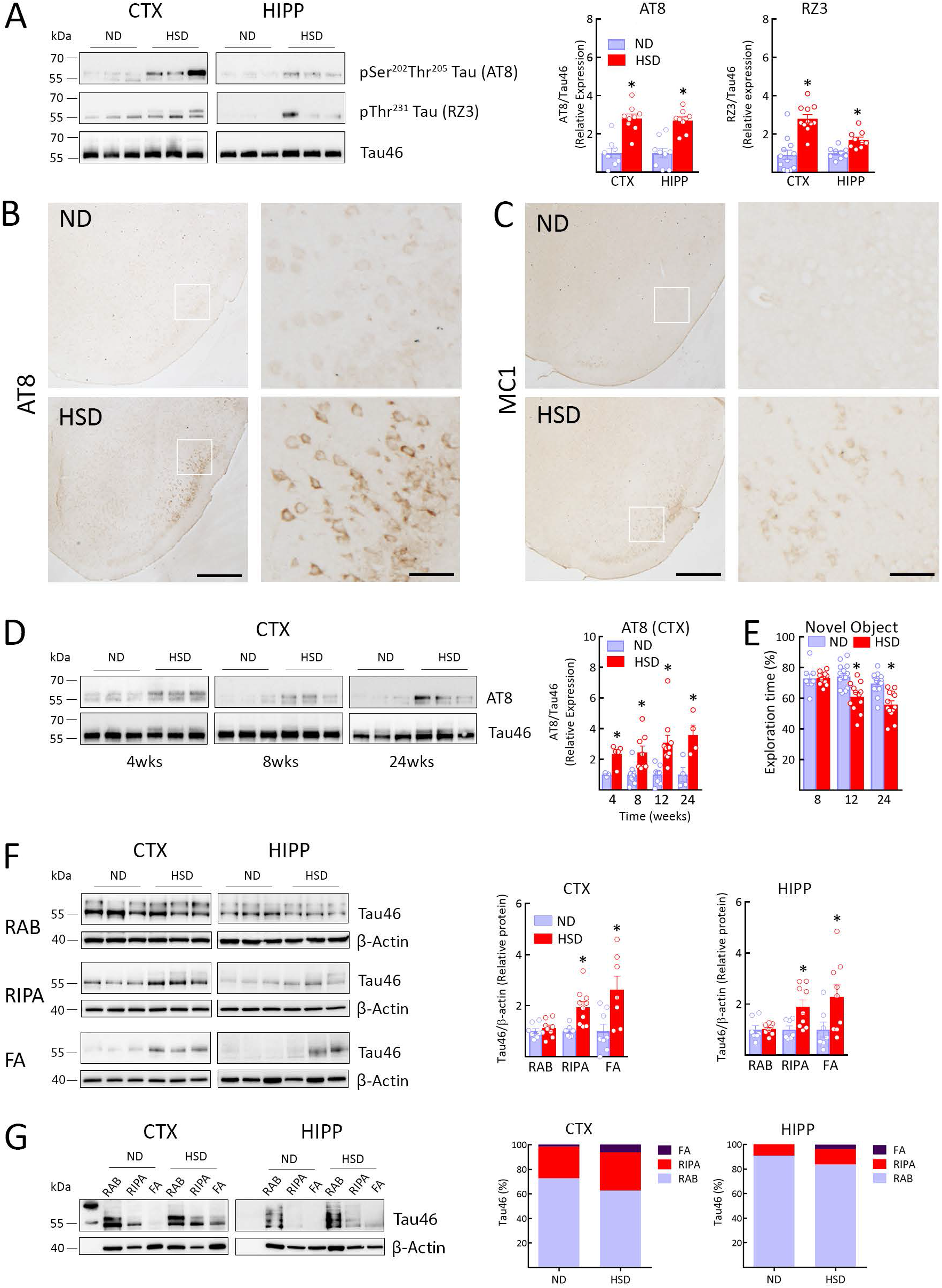
HSD increases tau phosphorylation and insoluble tau. A-B. HSD (NaCl 8%) increases tau phosphorylation on Ser^202^Thr^203^ (AT8) and Thp231 (RZ3) both in neocortex and hippocampus. (CTX: AT8, ND/HSD n=8/9, *p=0.0003 vs ND; RZ3, ND/HSD n=12/11, *p<0. 0001 vs ND; HIPP: AT8, ND/HSD n=9/9, *p=0. 0005 vs ND; RZ3, ND/HSD n=9/9, *p=0.0008 vs ND, two-tailed unpaired t-test). **C:** HSD increases AT8 and MC1 immunoreactivity in neuronal cell bodies of the piriform cortex (size bar=500pm; 100pm in inset). **D:** Time course of the increase in AT8 induced by HSD in the neocortex. The increase is observed at 4 weeks and is greatest at 24 weeks of HSD (AT8, 4 weeks: ND/HSD n=3/5, *p=0. 0357 vs ND 4wks; 8 weeks: ND/HSD n=9/8, *p=0.0016 vs ND 8wks; 24 weeks: ND/HSD n=4/4, *p=0. 0286 vs ND 24wks, two-tailed unpaired t-test). **E:** HSD induces deficits in recognition memory assessed by the novel object test, first observed at 12 weeks (Diet: *p<0.0001, Time: *p=0. 0002; 8 weeks: ND/HSD n=8/11; 12 weeks: ND/HSD n=16/12; 24 weeks: ND/HSD n=14/13 mice/group, two-way ANOVA and Tukey’s test). **F:** Extraction of total tau in RAB, RIPA and 70% FA buffer in neocortex and hippocampus. HSD increases levels of tau extracted in RIPA and FA after 12 weeks of treatment indicating increased insolubility (CTX: RIPA, ND/HSD n=7/10, *p=0. 0032 vs ND RIPA; FA, ND/HSD n=8/7, *p=0. 0146 vs ND FA; HIPP: RIPA, ND/HSD n=7/9, *p=0. 0186 vs ND RIPA; FA, ND/HSD n=7/9, *p=0.0494 vs ND FA, two-tailed unpaired t-test). **G:** Fractionation of total tau in RAB, RIPA, or 70% FA. HSD shifts tau from the more soluble RAB fraction to the less soluble RIPA and FA fractions (CTX: ND/HSD n=9/8, RAB, p=0.4234 vs ND, RIPA, p=0.5414 vs ND, FA, *p<0.0325 vs ND; HIPP: ND/HSD n=5/6, RAB, p=0.2468 vs ND, RIPA, p=0.3290 vs ND, FA, *p<0.0152 vs ND, two-tailed unpaired t-test). I mmunoblots in **A, D, F** and **G** are cropped. Full gel pictures for immunoblots are shown in Extended Data Fig.5 and 6. Data are expressed as mean±SEM.

In neocortex, the increase in AT8 was observed at 4 weeks of HSD and was greatest at 24 weeks, whereas in hippocampus p-tau peaked at 12 weeks and then declined (Fig. 1D and Extended Data Fig. 1G). An increase in RZ3 was observed at 8 and 12 weeks in neocortex and at 12 weeks in the hippocampus (Extended Data Fig. 1G). Starting at 12 weeks of HSD, mice exhibited difficulties in recognizing novel objects and developed a deficit in spatial memory at the Barnes maze, suggesting impaired cognition (Fig.1E; Extended Data Fig. 2A). Therefore, tau phosphorylation occurs in parallel with the endothelial NO deficit previously described^8^ and is followed by cognitive deficits. To determine if p-tau is upregulated also in other conditions associated with endothelial dysfunction, we investigated models of hypertension in which deficits in endothelial NO and cognitive function are well described^25^. We found an increase in p-tau in hypertension produced by chronic administration of the pressor peptide angiotensin-II or in BPH/2J mice with life-long elevations in blood pressure (Extended Data Fig. 2B-C).

**Figure 2.**
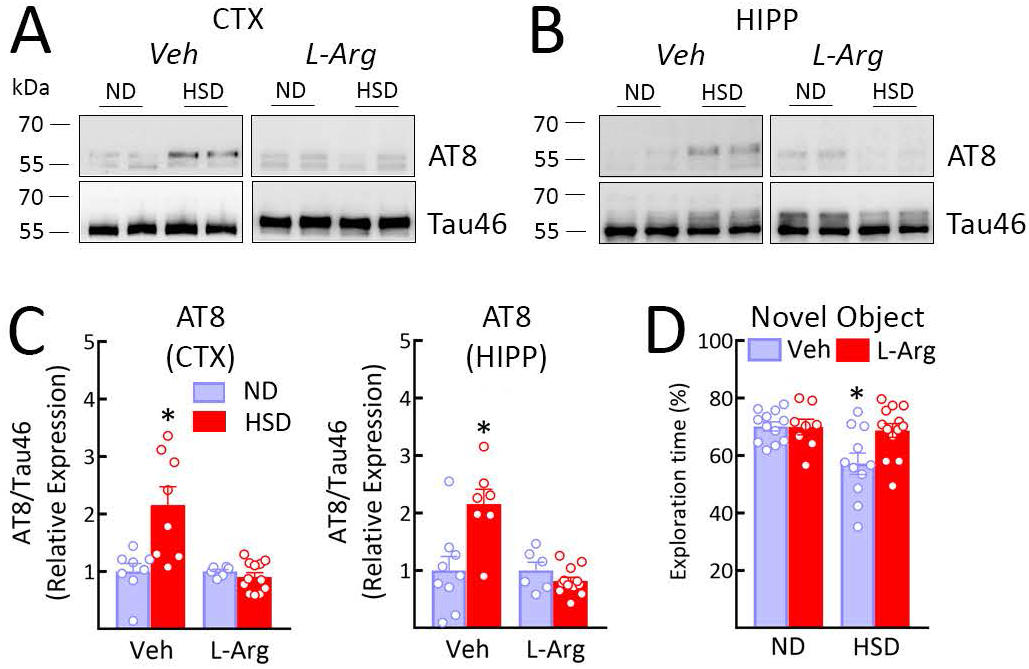
The NO precursor L-arginine prevents the increase in p-tau induced by HSD. A-C. Administration of L-arginine (10 g/L in drinking water), starting at week 8 of HSD and continued through week 12, suppresses AT8 accumulation in both neocortex and hippocampus (CTX: Vehicle, ND/HSD n=8/8, Diet: *p=0. 0060, Treatment: *p=0.0015; HIPP: Vehicle, ND/HSD n=9/7, Diet: *p=0.0145, Treatment: *p=0. 0011, two-way ANOVA and Bonferroni’s test). **D:** L-arginine treatment improves the cognitive deficits induced by HSD (Veh — ND/HSD n=12/10, L-Arg — ND/HSD n=6/11; Diet: *p=0. 0156, Treatment: *p=0. 0406, two-way ANOVA and Tukey’s test). lmmunoblots in **A** and **B** are cropped. Full gel pictures for immunoblots are shown in Extended Data Fig.7. Data are expressed as mean±SEM.

Tau solubility is a critical determinant of its harmful neuronal effects, and insoluble tau has been implicated in the neuronal dysfunction driving cognitive impairment^17^. To determine whether HSD alters tau solubility, tau levels were examined in neocortical and hippocampal lysates by Western blotting after sequential biochemical extraction in RAB (salt buffer), RIPA (detergent buffer) or 70% formic acid (FA), containing, respectively, soluble, less soluble and highly insoluble tau. After 12 weeks of HSD the tau in RIPA and FA fractions increased in neocortex and hippocampus, consistent with an increase in more insoluble tau (Fig. 1F-G). Since hypothermia in the setting of hibernation increases p-tau levels without causing cognitive impairment^26^, we performed a similar analysis in the brain of mice subjected to hypothermia. We found that hypothermia increases p-tau, but, at variance with HSD, does not produce a shift towards more insoluble species (Extended Data Fig. 2D-E). These observations indicate that HSD leads to tau phosphorylation and a shift from soluble to insoluble tau.

The NO precursor L-arginine counteracts the deficit in endothelial NO induced by HSD^8^ and other conditions associated with endothelial dysfunction^27,28^. Therefore, we asked if L-arginine would also rescue the p-tau accumulation induced by HSD and, if so, whether the effect is associated with improved cognition. To this end, mice were given L-arginine in the drinking water (10gr/L) during the last 4 weeks of the HSD or ND 12 week-treatment. We have previously demonstrated that L-arginine normalizes resting and stimulated cerebral endothelial NO synthesis without affecting arterial pressure^8^. L-arginine suppressed p-tau accumulation both in neocortex (AT8 and RZ3) and hippocampus (AT8) and prevented the cognitive dysfunction induced by HSD (Fig. 2A-D; Extended Data Fig. 3A).

**Figure 3.**
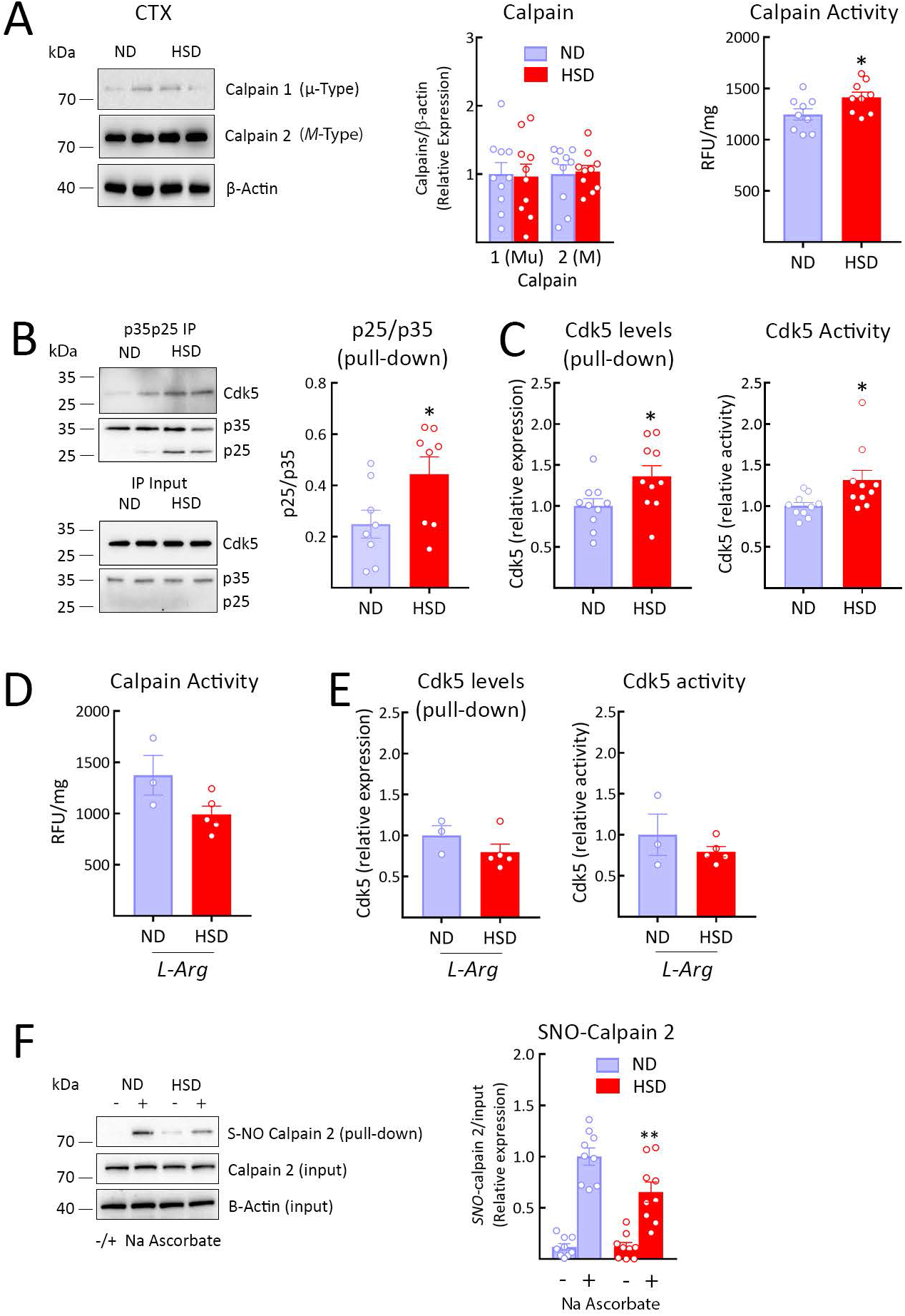
HSD induces activation of calpain and Cdk5, an effect associated with calpain denitrosylation. **A:** HSD did not alter calpain 1 or 2 expression (ND/HSD, n=10), but increased enzyme activity (ND/HSD n=9/9, *p=0.0314 vs ND, two-tailed unpaired t-test). **B:** Consistent with calpain activation, HSD increases the cleavage of p35 into p25 (ND/HSD n=8/8, *p=0. 0379 vs ND, two-tailed unpaired t-test). **C:** HSD increases the levels of Cdk5 bound to p35p25 (ND/HSD n=10/10, *p=0. 0232 vs ND, two-tailed unpaired t-test) and Cdk5 activity (ND/HSD n=10/10, *p=0.0115 vs ND, two-tailed unpaired t-test). **D-E:** L-arginine administration counteracts the increase in calpain and Cdk5 activity induced by HSD (ND/HSD n=3/5). **F:** Nitrosylation of calpain 2, the predominant brain isoform of the enzyme (see **A**), is reduced in HSD mice (ND/HSD n=9/9, Diet: *p=0.0189; Treatment: *p<0.0001, two-way ANOVA and Tukey’s test). Nitrosylation was assessed by the biotin switch assay. I mmunoblots in **A, B** and **F** are cropped. Full gel pictures for immunoblots are shown in Extended Data Fig.8. Data are expressed as mean±SEM.

Cdk5 is a major kinase responsible for tau hyperphosphorylation^29^. Cdk5 activity is tightly regulated by its protein binding partners, including p35^30^. In conditions associated with neuronal stress, cleavage of p35 into p25 by calpain leads to dysregulated activation of Cdk5 and hyperphosphorylation of its targets, including tau^31,32^. Since reduced endothelial NO may lead to tau phosphorylation by activating Cdk5 via p25^33^, we examined if HSD influences calpain and Cdk5 activity. Calpain 2 is more abundant than calpain 1 in neocortex (Fig. 3A), is located mainly in neurons (Extended Data Fig. 3B)^34^ and colocalizes with Cdk5 (Extended Data Fig. 3C). HSD did not alter calpain expression (Fig. 3A), but resulted in activation of the enzyme, leading to an increase in the p25/p35 ratio, in Cdk5 bound to p25/p35 and in Cdk5 catalytic activity (Fig. 3A-C). Attesting to the involvement of endothelial NO, L-arginine administration prevented the calpain activation induced by HSD and the resulting increase in the Cdk5/p25/p35 complex and enzyme activation (Fig. 3D-E). L-arginine did not alter calpain levels (Extended Data Fig. 3D). GSK3β has also been implicated in tau phosphorylation, but HSD did not increase the activity of this enzyme in neocortex (Extended Data Fig. 3E). Similarly, HSD did not alter the expression of the prolyl cis/trans isomerase Pin-1, a regulator of tau dephosphorylation^35^ (Extended Data Fig. 3F).

Next, we examined the potential mechanisms by which endothelial NO deficiency may influence calpain activity. Calpain, once activated by Ca^2+^, is regulated mainly by its endogenous inhibitor calpastatin and by nitrosylation by NO^36^, which suppress calpain activity^37^. Since HSD did not reduce calpastatin expression (Extended Data Fig. 3G), we used the biotin switch assay to investigate the effect of HSD on calpain nitrosylation. Consistent with the observed calpain activation, we found that HSD reduces calpain nitrosylation (Fig. 3F).

The findings thus far suggest that HSD leads to neuronal p-tau accumulation through a deficit in endothelial NO resulting in denitrosylation and activation of calpain, which, in turn increases p25 levels resulting in activation of Cdk5. However, HSD also lowers resting CBF and impairs the ability of endothelial cells to regulate CBF, which could contribute to impair cognition by reducing the delivery of oxygen and glucose to brain regions involved in cognitive function ^19,38^. Therefore, we examined the relative contribution of p-tau and neurovascular dysfunction to the cognitive deficits induced by HSD. We reasoned that if tau is critical for the cognitive dysfunction, then tau-null mice should be protected from the deleterious cognitive effects of HSD despite sustained cerebral hemodynamic dysfunction. Tau-null mice were placed on ND or HSD and cerebral endothelial vasomotor function and cognition were assessed 12 weeks later. As anticipated^39^, the performance of tau-null mice to the novel object recognition test and Barnes maze was not different from that of WT controls fed a ND (Fig. 4A-C). Tau-null mice on HSD did not exhibit cognitive impairment, but still exhibited marked endothelial dysfunction, as reflected by the suppression of the CBF increase induced by bathing the neocortex with acetylcholine (Fig. 4D), a prototypical endothelial response mediated by eNOS-derived NO^38^. Therefore, it would seem that CBF dysregulation is not required for the cognitive dysfunction of HSD. To provide further evidence in support of this conclusion we treated WT mice with anti-tau antibodies (HJ8.8) or control IgG (50 mg/kg/week; i.p.) for the last 4 weeks of the 12-week ND or HSD regimen^40^. Anti-tau antibodies were previously shown to ameliorate cognitive function in a tauopathy mouse model^40^. In HJ8.8-treated mice, HSD induced a reduction in resting CBF and an attenuation of the endothelial CBF response to acetylcholine comparable to that observed in HSD-fed mice treated with control IgG (Fig. 4E-F). Despite persistent hypoperfusion and endothelial dysfunction, anti-tau antibodies ameliorated the cognitive deficit induced by HSD (Fig. 4G). The effect was associated with a reduction in p-tau in neocortex (AT8) and hippocampus (AT8 and RZ3) (Fig. 4H). As before^8^, HSD did not affect the increases in CBF induced by neural activity (Extended Data Fig. 4A)

**Figure 4.**
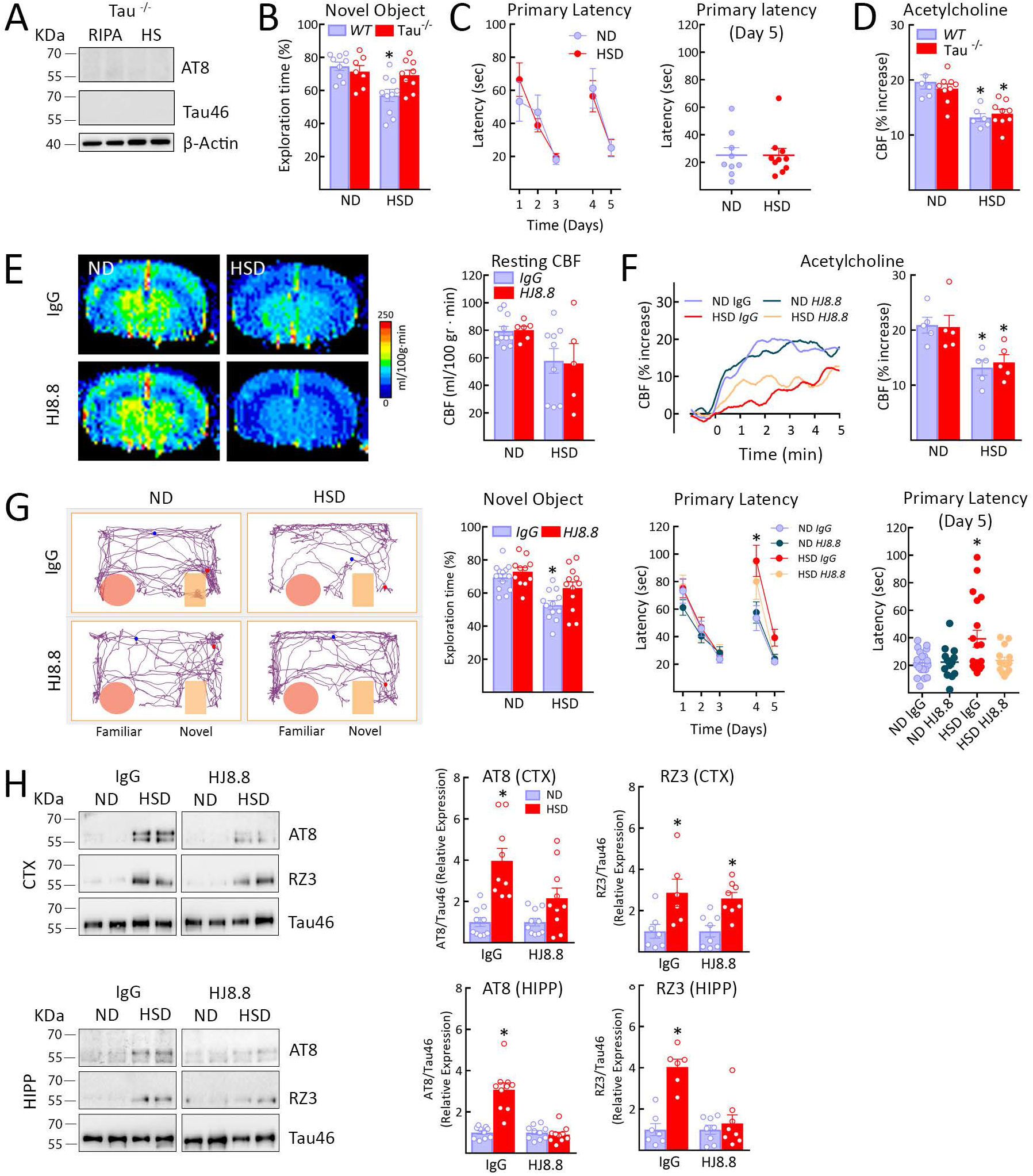
HSD-induced cognitive dysfunction is not observed in tau^-/-^ mice and prevented by tau antibodies despite cerebrovascular insufficiency. **A:** AT8 and Tau46 are absent in tau^-/-^ mice in both the RIPA and heat-stable RIPA fractions. **B-C:** HSD does not alter cognitive function in tau^-/-^ mice, assessed by the novel object recognition test (WT: ND/HSD n=9/10; Tau-/-: ND/HSD n=7/9; Diet: *p=0. 0055, Genotype: p=0.1827, two-way ANOVA and Tukey’s test) or the Barnes maze (ND/HSD n=9/10, Diet: p=0.9348, Time: *p<0001, Two-way ANOVA and Tukey’s test). **D:** Despite improved cognition, the CBF increase produced by neocortical application of acetylcholine in anesthetized mice equipped with a cranial widow, a response mediated by eNOS-derived NO, is still reduced in tau^-/-^ mice (ND/HSD WT, n=6/6, Tau-/-, n=9/9; Diet: *p<0. 0001, Genotype: p=0.7920, Two-way ANOVA and Tukey’s test). CBF was measured by laser-Doppler flowmetry. **E:** Systemic administration of anti-tau antibodies (HJ8. 8, 50mg/Kg/week i.p.) does not rescue the reduction in resting CBF induced by HSD, assessed by ASL-MRI (lgG: ND/HSD n=11/9; HJ8.8: ND/HSD n=6/5; Diet: *p=0.0061, Treatment: p=0.9367, two-way ANOVA and Tukey’s test). **F:** Similarly, HJ8.8 administration does not prevent the attenuation of the CBF response to ACh induced by HSD (lgG: ND/HSD n=5/5; HJ8.8: ND/HSD n=5/5; Diet: *p=0. 0005, Treatment: p=0.8516, twoway ANOVA and Tukey’s test). **G:** HJ8.8 administration ameliorates the cognitive dysfunction induced by HSD both at the novel object test (lgG: ND/HSD n=13/12; HJ8.8: ND/HSD n=11/12; Diet: *p<0.0001, Treatment: *p=0.0231, two-way ANOVA and Tukey’s test) and the Barnes maze (Primary Latency - lgG: ND/HSD n=19/19; HJ8.8: ND/HSD n=13/14; Time: *p<0. 0061, Diet: *p=0. 0317, repeated-measures two-way ANOVA and Tukey’s test; Primary Latency - Day 5 - lgG: ND/HSD n=19/19; HJ8.8: ND/HSD n=13/14; Diet: *p=0.0320, Treatment: p=0.0756, two-way ANOVA and Tukey’s test). **H:** HJ8.8 administration reduces AT8 levels in both neocortex and hippo-campus (AT8, CTX - lgG: ND/HSD n=10/9; HJ8.8: ND/HSD n=10/10; Diet: *p<0.0001, Treatment: *p=0. 0313; AT8 — HIPP - lgG: ND/HSD n=10/10; HJ8.8: ND/HSD n=10/10; Diet: *p<0.0001, Treatment: *p<0.0001, two-way ANOVA and Bonferroni’s test) and RZ3 levels in the hippocampus but not neocortex (RZ3, CTX - lgG: ND/HSD n=7/7; HJ8.8: ND/HSD n=8/8; Diet: *p=0.0005, Treatment: p=0.9586, RZ3 — HIPP - lgG: ND/HSD n=7/6; HJ8.8: ND/HSD n=8/8; Diet: *p<0. 0001, Treatment: *p=0.0004, two-way ANOVA and Bonferroni’s test). I mmunoblots in **A** and **H** are cropped. Full gel pictures for immunoblots are shown in Extended Data Fig.9. Data are expressed as mean±SEM.

These observations, collectively, are consistent with the hypothesis that the reduction in endothelial NO induced by HSD leads to calpain denitrosylation resulting in activation of the enzyme. The ensuing increase in p25 activates Cdk5 leading to tau phosphorylation in neurons, which, in turn, is responsible for the cognitive impairment (Extended Data Fig. 4C). In addition, the data indicate that the neurovascular dysfunction associated with HSD, also caused by the NO deficit, is not critical for the cognitive dysfunction. This conclusion is supported by the observations that the cognitive deficits induced by HSD do not occur in tau-null mice and are ameliorated by anti-tau antibodies, despite altered endothelium-dependent vasodilatation and reduced CBF.

The findings provide novel insights into the mechanisms by which cerebrovascular dysfunction alters cognition. Although to be confirmed in other cerebrovascular risk factors, the CBF reduction and suppression of endothelium-dependent vasoreactivity associated with HSD do not drive cognitive impairment. Whereas the suppression in endothelial NO induced by HSD is required for the cognitive impairment, the hemodynamic consequences of such NO deficit do not play a role. Rather, the cognitive dysfunction is dependent on the tau phosphorylation promoted by the deficit in endothelial NO. Thus, the cerebral hypoperfusion resulting from NO deficiency seems to be inconsequential to the cognitive deficit. Other aspects of endothelial function are at play, namely endothelial NO maintaining calpain homeostasis and preventing Cdk5 dysregulation and tau hyperphosphorylation.

Our data also provide a previously-unrecognized link between dietary habits, vascular dysfunction and tau pathology, independently of cerebral hypoperfusion. Such relationship may play a role in the frequent overlap between vascular and neurogenerative pathologies underlying late-life dementia ^41^. Whereas avoiding excessive salt consumption may help prevent tau pathology, therapeutic efforts to counteract cerebrovascular dysfunction need to go beyond rescuing cerebral perfusion, and target vascular mediators governing neurovascular interactions critical for cognitive health.

## MATERIALS AND METHODS

Most of the methods used in this study are well established in the laboratory and have been described in detail in previous publications^8,25,42^. Here we provide only a brief description.

### Mice

All procedures are approved by the institutional animal care and use committee of Weill Cornell Medicine (Animal protocol number: 0807-777A). Studies were conducted, according to the ARRIVE guidelines (https://www.nc3rs.org.uk/arrive-guidelines), in the following lines of mice: C57BL/6 (JAX), B6.129X1-Mapttm1Hnd (Tau ^-/-^, JAX, Stock #007251) and Tg(Camk2atTA)1Mmay Fgf^14Tg(tetO-MAPT*P301L)4510Kha/J^ (rTg4510, JAX, Stock#024854), BPH/2J mice (Age: 5 months; JAX, Stock #003005) and BPN/3J mice (Age: 5 months; JAX, Stock #003004). Unless otherwise indicated, male mice were used.

### High Salt Diet

Male or female mice (8 weeks old) received normal chow (0.5% NaCl) and tap water ad libitum (normal diet) or sodium-rich chow (4-8% NaCl) and tap water containing 1% NaCl ad libitum (HSD) for 4 to 24 weeks according to the experiment^8^.

### In vivo treatments

The nitric oxide precursor, L-arginine (10gr/L; Sigma) was administered in the drinking water starting at 8 weeks of HSD and continuing until 12 weeks. ND and HSD mice were treated (i.p., weekly) with 50mg/kg of anti-Tau (HJ8.8) or mouse IgG1 isotype control (Clone MOPC-21; bioXcell) antibodies for the last 4 weeks of the HSD treatment period (12 weeks) prior to behavioral and cerebrovascular studies.

### General surgical procedures for CBF studies

Mice were anesthetized with isoflurane (induction, 5%; maintenance, 2%). The trachea was intubated and mice were artificially ventilated with a mixture of N_2_ and O_2_. One of the femoral arteries was cannulated for recording mean arterial pressure (MAP) and collecting blood samples for blood gas analysis^43^. Rectal temperature was maintained at 37°C. End tidal CO_2_, monitored by a CO_2_ analyzer (Capstar-100, CWE Inc.), was maintained at 2.6–2.7% to provide a pCO_2_ of 30–40 mmHg and a pH of 7.3-7.437. After surgery, isoflurane was discontinued and anesthesia was maintained with urethane (750 mg/kg, i.p.) and chloralose (50 mg/kg, i.p.). Throughout the experiment the level of anesthesia was monitored by testing motor responses to tail pinch.

### Monitoring cerebral blood flow

A small craniotomy (2×2mm) was performed to expose the parietal cortex, the dura was removed, and the site was superfused with Ringer’s solution (37°C; pH 7.3–7.4)^25^. CBF was continuously monitored at the site of superfusion with a laser-Doppler probe (Perimed) positioned stereotaxically ≈0.5mm above the cortical surface and connected to a data acquisition system (PowerLab). CBF values are expressed as percentage increases relative to the resting level.

### Protocol for CBF experiments

After MAP and blood gases stabilized, CBF responses were recorded^8^. The whisker-barrel cortex was activated for 60 seconds by stroking the contralateral vibrissae, and the evoked changes in CBF were recorded. The endothelium-dependent vasodilator acetylcholine (ACh; 100µM; Sigma), was superfused on the exposed neocortex for 5 minutes and the associated CBF changes were recorded by laser-Doppler flowmetry.

### Measurement of resting CBF by ASL-MRI

CBF was assessed quantitatively using arterial spin labeling magnetic resonance imaging (ASLMRI) as previously described^8^. The ASL images were analyzed by ImageJ and the average CBF value is reported as mL per 100g of tissue per minute.

### Osmotic minipumps implantation

Osmotic minipumps containing vehicle (saline) or ANG II (600 ng·kg−1·min−1) were implanted subcutaneously under isoflurane anesthesia. Systolic blood pressure was monitored in awake mice using tail-cuff plethysmography^25^. Forty-two days later, mice were anesthetized and their brains were collected for assessment of tau phosphorylation.

### Hypothermia

C57BL/6 mice (12 weeks old) were anesthetized by injection of ketamine/xylazine (100/10 mg/kg). Rectal temperature was continuously monitored and kept at 37°C (normothermia) or 30°C (hypothermia) using a thermostatically-controlled heating pad. Mice were sacrificed 30 minutes after anesthesia and their brains were collected and frozen on dry ice. Tissues were kept at −80°C until processing for immunoblot analysis.

### Immunoblot analysis

Cortex (≈80-90mg) and hippocampus (≈15mg) isolated from ND and HSD mice were sonicated in 800 and 600μl of RIPA buffer (50mM Tris-HCl pH 8.0, 150mM NaCl, 0.5% Deoxycholic Acid, 0.1% SDS, 1mM EDTA pH 8.0, 1% IGEPAL CA-630, 1mM Na3VO4, 20mM NaF and one tablet/10mL of cOmplete™, EDTA-free Protease Inhibitor Cocktail, Millipore Sigma) and equal volumes were mixed with SDS sample buffer, boiled, and analyzed on 10% or 10-20% Novex™ WedgeWell™ gels (Thermo Fisher Scientific). Proteins were transferred to PVDF membranes (Millipore), blocked at room temperature (RT) for 1 hour with 5% milk in TBS, and incubated, overnight at 4°C, with primary antibodies (see Extended Data Table 1) in 5% BSA in TBS/0.1% Tween-20 (TBST). Membranes were washed in TBST, incubated with goat anti-mouse or rabbit secondary antibodies conjugated to horseradish peroxidase (Santa Cruz Biotechnology) for 1 hour at RT and protein bands were visualized with Clarity Western ECL Substrate (Bio Rad) on a Bio Rad ChemiDoc MP Imaging System. Quantification was performed using Image Lab 6.0 (Bio Rad).

### Heat-stable fractions

After homogenization in cold RIPA buffer and centrifugation, 150µl of the supernatant containing the proteins was boiled at 100°C for 10 minutes. Samples were cooled on ice for 20 minutes and then centrifuged at 20,000 g at 4°C for 15 minutes. The supernatant corresponding to the heat stable (HS) fraction was then harvested. This method is used to isolate proteins resistant to heat including tau and other microtubule-associated proteins (MAPs). Thus, endogenous immunoglobulins are precipitated during the boiling process and eliminated from the supernatant. The proteins were then mixed with equal volumes of SDS sample buffer, boiled, and analyzed on 10% Novex™ WedgeWell™ gels (Thermo Fisher Scientific). Although Tau protein is partially lost during the boiling process, the HS samples are enriched with Tau (please see Extended Data Fig. 4B). Furthermore, boiling significantly improves specificity of certain antibodies such as AT8, RZ3 or MC1^44^.

### Tau dephosphorylation

After overnight dialysis to remove phosphatase inhibitors, protein samples (40µl) were incubated with 5μL of 10X NEBuffer for Protein MetalloPhosphatases (PMP), 5μL of 10mM MnCl_2_ and 1µl of Lambda Protein Phosphatase (Lambda PP, New England Biolabs) at 30°C for 3 hours. Reactions were stopped by addition of SDS sample buffer and boiling for 5 minutes at 100 °C.

### Brain tissue protein extraction

Extraction was performed as described previously^40^. The cortex (≈80-90mg) and the hippocampus (≈15mg) of each brain were homogenized by sonication in 800 and 300μl of RAB buffer [100mM MES, 1mM EDTA, 0.5mM MgSO_4_, 750mM NaCl, 20mM NaF, 1mM Na3VO4, supplemented by EDTA-free Protease Inhibitor Cocktail, Millipore Sigma], respectively. In brief, the samples were centrifuged at 50,000g for 20 minutes at 4°C using an Optima MAX-TLA 120.2 Ultracentrifuge (Beckman). The supernatants were collected as RAB soluble fractions and pellets were resuspended in identical volumes of RIPA buffer [150mM NaCl, 50mM Tris, 0.5% deoxycholic acid, 1% Triton X-100, 0.5% SDS, 25mM EDTA, pH 8.0, 20mM NaF, 1mM Na_3_VO_4_ supplemented by EDTA-free Protease Inhibitor Cocktail, Millipore Sigma], and centrifuged at 50,000 g for 20 minutes at 4°C. The supernatants were collected as RIPA soluble fractions. The pellets were sonicated in 70% formic acid (300μl for the cortex and 125μl for the hippocampus), and centrifuged at 50,000g for 20 minutes at 4°C. The supernatants were collected as 70% formic acid fractions. All fractions were stored in −80°C until analyzed. For western blotting, an aliquot of 100μl of the formic acid fractions was evaporated in a Savant SpeedVac concentrator at 45°C for 1 hour. The samples were resuspended in 100μl of SDS sample buffer with the addition of 1μl of 10N NaOH, sonicated and then boiled for 5 minutes.

### Immunohistochemistry

After 12 weeks of ND/HSD, mice were anesthetized with intraperitoneal pentobarbital (200 mg/kg), and then perfused transcardiacally with cold PBS, followed by cold 4% paraformaldehyde (PFA) in PBS. The brains were removed and immersed first in 4% PFA overnight and then in 70% ethanol for 3 days. Brains were then embedded in paraffin and cut into 6μm sections using a microtome. After rehydration and antigen retrieval in preheated citrate buffer (10μM) for 30 minutes, brain sections were immersed in 3% H_2_O_2_ and then blocked with 100% Sniper (Biocare Medical) for 1 hour. After blocking, sections were incubated for 2.5 days at 4°C with the AT8, MC1, Calpain 2 or Cdk5 antibody (1:250, 1:100, 1:100 and 1:100 in 1:50 Sniper in PBS, respectively) and thereafter processed for 1 hour with the biotinylated secondary antibody in 1% normal donkey serum PBS (anti-mouse IgG1, Jackson ImmunoResearch) or Cy3 anti-rabbit and FITC anti-mouse (Jackson ImmunoResearch) for immunofluorescence studies. Reactions were visualized with the ABC-complex (Vectorlabs) and 3,3-diaminobenzidine. A Nikon light microscope or a confocal microscope (Leica TCS SP5) was used to visualize the signal associated with each antibody.

### Thioflavin S Staining

After mounting on slides and post-fixation with 4% PFA in PBS for 10 minutes, coronal brain sections (40µm) were washed and labeled with 0.05% (wt/vol) thioflavine-S in 50% (vol/vol) ethanol for 10 minutes as previously described^45^. An epifluorescence microscope (IX83 Inverted Microscope, Olympus) was used to visualize the FITC signal associated with thioflavine-S.

### Calpain activity

Calpain activity was measured by using a Calpain Activity Assay Kit from AbCam^46,47^. Briefly, fresh cortex and hippocampus were homogenized in the extraction buffer provided with the kit, which specifically extracts cytosolic proteins without contaminations of cell membrane and lysosome proteases and prevents auto-activation of calpain during the extraction procedure. The fluorometric assay is based on the detection of cleavage of calpain substrate Ac-LLY-AFC. AcLLY-AFC emits blue light (ʎmax = 400nm); upon cleavage of the substrate by calpain, free AFC emits a yellow-green fluorescence (ʎmax = 505nm), which can be quantified using a fluorometer or a fluorescence plate reader. Specificity of the signal was confirmed by using the calpain inhibitor Z-LLY-FMK (100-200µM). The activity is expressed as Relative Fluorescent Unit (RFU) per milligram of protein for each sample.

### p35/p25 and GSK3β immunoprecipitation

Immunoprecipitation was performed with anti-p35p25 (Cell Signaling), anti-GSK3β (Cell Signaling) or anti-rabbit monoclonal IgG1 isotype control antibody (Santa Cruz Biotechnology). Samples were incubated overnight with the primary antibodies and then with protein-A sepharose (p35p25) (GE Healthcare Life Sciences) or protein-G Dynabeads (GSK3β) (Thermo Fisher Scientific) for 2 hours at 4º C. Precipitates were used for Cdk5 or GSK3β activity measurements. Immunoprecipitation was confirmed by loading the samples on 10% Tris-glycine SDS polyacrylamide gels and western blot as described above.

### Detection of S-nitrosylation of calpain 2 with the biotin-switch technique

Detection of S-nitrosylated calpain 2 was performed using the biotin-switch technique, as previously described^48^. Briefly, samples were sonicated in 800μl of RIPA buffer containing 0.1mM of neocuproine and, after centrifugation, protein concentrations were measured. Cysteine thiol groups in 1mg of proteins were blocked with 10% S-methylmethane thiosulfonate (MMTS) (Sigma). After protein-precipitation with 100% acetone, sodium ascorbate was added to the sample to convert each S-nitrothiols (SNO) to a free thiol via a transnitrosation reaction to generate O-nitrosoascorbate. Next, each nascent free thiol (previously an SNO site) was biotinylated with biotin–HPDP (Pierce). Biotinylated proteins were then pull-down by using avidin beads and analyzed on 10% Novex™ WedgeWell™ gels (Thermo Fisher Scientific). Before avidin pulldown, a small fraction of each sample was collected to determine protein “input.” The degree of pulldown correlates with protein S-nitrosylation of calpain 2 which was detected with an antibody against the protein (see Extended Data Suppl. Table 1). Nitrosylation of calpain 2 is expressed as the ratio between the pull-down signal and the input corrected for the β-actin levels.

### Cdk5 and GSK3β activity

Cdk5 activity in brain lysates was determined after pulldown with p25/p35 antibody (Cell Signaling) from 500µg total protein using a synthetic histone H1 peptide substrate (PKTPKKAKKL, Enzo Life Sciences). GSK3β activity was determined after pulldown with GSK3β antibody (Cell Signaling) from 100µg total protein using phospho-glycogen synthase peptide-2a as substrate (Tocris). Phosphorylation reactions were initiated by mixing bead-coupled Cdk5 with 40µl reaction buffer containing the following: 50mM HEPES.KOH (pH 7.4), 5mM MgCl_2_, 0.05% BSA, 50µM substrate, 50µM cold ATP, 1mM dithiothreitol, 1x complete protease inhibitors without EDTA (Roche Applied Biosciences) and 5 Ci/mmole ^γ32^P-ATP. Companion reactions for every sample were executed in the presence of the Cdk5 inhibitor ((R)-CR8, Tocris) (10µM) or the GSK3β inhibitor (CHIR 99021, Tocris) (10µM) to correct for non-specific activity. Reactions were incubated at 30°C for 30 minutes, after which they were terminated by spotting on P81 phosphocellulose cation exchange chromatography paper. Filters were washed 4 times for 2 minutes in 0.5% phosphoric acid, and the remaining radioactivity was quantified in a scintillation counter by the Cherenkov method.

### Novel Object Recognition Test

The novel object recognition test (NOR) task was conducted under dim light in a plastic box. Stimuli consisted of plastic objects that varied in color and shape but had similar size^49,50^. A video camera mounted on the wall directly above the box was used to record the testing session for off-line analysis. Mice were acclimated to the testing room and chamber for one day prior to testing. Twenty-four hours after habituation, mice were placed in the same box in the presence of two identical sample objects and were allowed to explore for 5 minutes. After an intersession interval of 1 hour, mice were placed in the same box but one of the two objects was replaced by a novel object. Mice were allowed to explore for 5 minutes. Exploratory behavior was later assessed manually by an experimenter blinded to the treatment group. Exploration of an object was defined as the mouse sniffing the object or touching the object while looking at it. Placing the forepaws on the objects was considered as exploratory behavior but climbing on the objects was not. A minimal exploration time for both objects (total exploration time) during the test phase (~5 seconds) was used. The amount of time taken to explore the novel object was expressed as percentage of the total exploration time and provides an index of recognition memory^49,50^.

### The Barnes Maze test

The Barnes maze consisted of a circular open surface (90cm in diameter) elevated to 90cm by four wooden legs^51^. There were 20 circular holes (5cm in diameter) equally spaced around the perimeter, and positioned 2.5cm from the edge of the maze. No wall and no intra-maze visual cues were placed around the edge. A wooden plastic escape box (11×6×5cm) was positioned beneath one of the holes. Two neon lamps and a buzzer were used as aversive stimuli. The Any-Maze tracking system (Stoelting) was used to record the movement of mice on the maze. Extra-maze visual cues consisted of objects within the room (table, computer, sink, door, etc.) and the experimenter. Mice were tested in groups of seven to ten, and between trials they were placed into cages, which were placed in a dark room adjacent to the test room for the inter-trial interval (20-30 minutes). No habituation trial was performed. The acquisition phase consisted of 3 consecutive training days with three trials per day with the escape hole located at the same location across trials and days. On each trial a mouse was placed into a start tube located in the center of the maze, the start tube was raised, and the buzzer was turned on until the mouse entered the escape hole. After each trial, mice remained in the escape box for 60 seconds before being returned to their cage. Between trials the maze floor was cleaned with 10% ethanol in water to minimize olfactory cues. For each trial mice were given 3 minutes to locate the escape hole, after which they were guided to the escape hole or placed directly into the escape box if they failed to enter the escape hole. Four parameters of learning performance were recorded: (1) the latency to locate (primary latency) and (2) enter the escape hole (total latency), (3) the number of errors made and (4) the distance traveled before locating the escape hole^51^. When a mouse dipped its head into a hole that did not provide escape was considered an error. On days 4 and 5, the location of the escape hole was moved 180° from its previous location (reverse learning) and two trials per day were performed.

### Statistics

Sample size was determined according to power analysis based on previously published work by our lab on the effects of dietary salt on CBF regulation and cognitive function. On these bases, 10-15 mice/group were required in studies involving assessment of cognitive function and cerebrovascular function^8,25^. Mouse randomization was performed based on the random number generator function (RANDBETWEEN) in Microsoft Excel software. Analysis of the data was performed in a blinded fashion and GraphPad Prism (v. 6.0) software was used for statistical analysis. Intergroup differences were analyzed by unpaired Student’s t-test for single comparison or by one or two-way analysis of variance (Tukey’s or Bonferroni’s post-hoc analysis) for multiple comparisons. Data are expressed as mean±SEM and differences are considered statistically significant for p<0.05.

## Data Availability

All data generated or analyzed during this study are included in this published article (and its supplementary information files).

## ACKNOWLEDGEMENTS

We thank Prof. Peter Davies for providing the RZ3, MC1 and PHF1 antibodies. This study was supported by National Institutes of Health grants R37-NS089323 (CI) and 1R01-NS095441 (CI), by a grant from the Cure Alzheimer’s Fund (GF and CI), and by a Scientist Development Grant from the American Heart Association (GF). The support from the Feil Family Foundation is gratefully acknowledged.

## AUTHOR CONTRIBUTIONS

G.F. performed western blotting experiments, behavioral tests, cerebrovascular studies and analyzed data. K.H. performed experiments on Cdk5/GSK3β activity and analyzed data. S.G.S. performed western blotting experiments, behavioral tests and immunohistochemistry. S.S. and M.M.S. performed experiments on the effects of hypertension on tau. A.M. performed immunohistochemistry experiments. H.J. and D.M.H. provided the HJ8.8 antibody. J. A supervised the molecular aspects of the study and edited the manuscript. G.F. and C.I. designed and supervised the entire study and wrote the manuscript.

## COMPETING INTERESTS

D.M.H. is listed as an inventor on a patent licensed by Washington University to C2N Diagnostics and subsequently AbbVie on the therapeutic use of anti-tau antibodies. D.M.H. co-founded and is on the scientific advisory board of C2N Diagnostics. D.M.H. is on the scientific advisory board of Denali, Genentech, and Proclara.

